# Corrective feedback control of competing neural network with entire connections

**DOI:** 10.1101/2022.01.10.475737

**Authors:** Uramogi Wang

## Abstract

Continuous persist activity of the competitive network is related to many functions, such as working memory, oculomotor integrator and decision making. Many competition models with mutual inhibition structures achieve activity maintenance via positive feedback, which requires meticulous fine tuning of the network parameters strictly. Negative derivative feedback, according to recent research, might represent a novel mechanism for sustaining neural activity that is more resistant to multiple neural perturbations than positive feedback. Many classic models with only mutual inhibition structure are not capable of providing negative derivative feedback because double-inhibition acts as a positive feedback loop, and lack of negative feedback loop that is indispensable for negative derivative feedback. Here in the proposal, we aim to derive a new competition network with negative derivative feedback. The network is made up of two symmetric pairs of EI populations that the four population are completely connected. We conclude that the negative derivative occurs in two circumstances, in which one the activity of the two sides is synchronous but push-pull-like in the other, as well as the switch of two conditions in mathematical analysis and numerical simulation.

## Introduction

Many cortical functions, such as parametric working memory, decision making and neural integrator, are thought to be substantially tied to the persistent neural activity induced by the competitive neural network^[1]^. The traditional competitive structure is made up of two inhibitory populations connected by mutual inhibition, which is required for competitive activities. To explain working memory in two-interval discriminating choice tasks, Machens and Romo devised a basic model without self -connections to each side^[2]^. For oculomotor control and the neural integrator, several studies employ a similar paradigm^[3]^. However, according to Dale’s law, two mutual inhibitory populations don’t have self-excitation connections, which is ubiquitous in experimental recordings. Therefore, the structure of two mutual inhibition network is insufficiently physiologically rational.

X-J Wang has presented a more biologically sensible competition model^[4]^. In the model,7200 LIF(Leaky Integrate and Fire) spiking neurons compose a three-population neural architecture network, in which two excitatory neuron populations inhibit each other through a common inhibitory neuron pool. The emphasis of the study is on discrete attractor analysis of the decision-making process, rather than persistent activity. Furthermore, all these models show that the only way to sustain persistent neural activity is via positive feedback mechanisms^[5]^. Generally speaking, most mutual inhibition structure composes a double-negative connection loop that is equivalent to a positive feedback pathway. Positive feedback inputs might be used to balance the intrinsic leakage activity to maintain the information^[6]^.However, positive feedback is supposed to have precise network parameter, requires a fine tuning of the level of feedback and are extremely susceptible to common perturbations, such as global changes in neuronal or synaptic excitabilities, that disrupt this tuning.

In the brain, the representation and processing of information have great temporal and spatial dynamics. Furthermore, the presence of noise and physical changes (decline of neurons, change of synaptic strength, etc.) leads to the conclusion that network should be robust against common perturbations to some extent. Therefore, Subkin Lim and Mark.S Goldman^[7]^ introduce corrective feedback control into cortical microcircuit. The structure is made up of one excitatory and one inhibitory population with recurrent connections, consisting of positive feedback pathway and negative feedback pathway in which the connection strengths are equal, but distinct synaptic constants. This accords with the principle of corrective input in control theory: negative derivative feedback arises when recurring excitatory and inhibitory inputs to the population are balanced in strength and offset in time.

We construct a four-population network consisting of two pairs of symmetric EI populations to apply negative derivative feedback to competition network. The structure is divided into three parts. First, the reason why the number of populations is four, with two excitatory and two inhibitory populations. Because Dale’s law states that if two neurons have inhibitory connections, they can only be inhibitory neurons. In traditional two-population network with mutual inhibition, the neurons can only be inhibitory and it restricts excitatory connections and activities and is biologically unreasonable. Secondly, the reason why the populations are completely connected. Mutual inhibition may be established in a four-population structure by activating contralateral inhibitory neurons or inhibition of contralateral excitatory neurons (E to I cross and I to E cross). However, if there are no alternative negative feedback routes, simply mutual inhibition connections are not capable of negative derivative feedback. How to achieve those negative feedback pathways? In fact, it is really a complex problem. Therefore, we treat the model as if it were a fully connected network, the most complicated instance with both positive and negative feedback pathways originally. Furthermore, since cortical neurons receive vast quantities of both excitation and inhibition in a broad variety of situations and brain locations, a comprehensive connection is feasible for biological consideration^[8]^.It is also an ideal situation as a mathematical model.

Finally, however, the difficulty is to derive negative derivative feedback from the structure briefly. Due to the complexity of the network structure, pathways of positive feedback and negative feedback intersect within each other, which leads to obscure differentiation and definition of positive or negative feedback pathways. The two EI populations are assumed symmetric for the sake of dimension reduction. Another reason is push-pull like activity needs a symmetric structure to keep the equivalent role of two sides.

The underlying mechanism by which negative derivative feedback might lead to persistent activity in the linked four population structure was investigated. Our analysis is based on the assumption that symmetrical network structure leads to symmetrical activities without any input. Therefore, when the system reaches stability, the activities on both sides are always the same or “opposite” compared with background baseline activity, resulting in two persistent activity modes in which one the activity of the two sides is synchronous but push-pull-like in the other. Finally, using mathematical analysis and numerical simulation, we were able to determine the assumption and infer the transition between two modes.

## Method

The model is a four-population network consists of two same excitatory populations and two inhibitory populations. Connection strengths and time constants are symmetric distributed in two sides. The structure of the network is shown as:

As shown in the figure, each population connects with the all other populations. *EE, EI, IE, II* represent the connections within each side, and, *EEC, EIC, IEC, IIC* represent the connections from one side to the other side. We denote connection strengths and time constants as *J*_*i*_ and *τ*_*i*_ with *i* ∈ {*EE, EI, IE, II, EEC, EIC, IEC, IIC*}. The firing rates of the four population are denoted as *R*_*Ej*_ and *R*_*Ij*_ with *j* ∈ {1,2}. The synaptic activities of the connections are denoted as *S*_*ij*_. The system is described by linear firing rate model as:

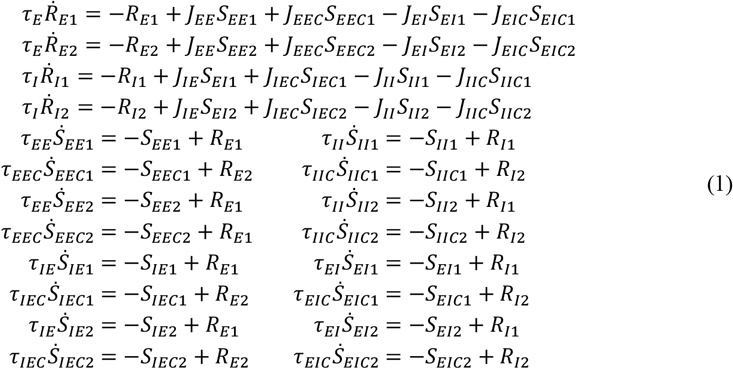

The system above could be described as 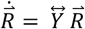, of which the vector 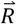 represents the activity of neural population and 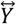 represents the matrix of network parameters. To solve the system, we have

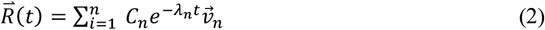

where λ_*n*_ and 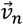 are the eigenvalue and eigenvector of the matrix 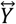 with n dimensions. For the require of persist activity, at least one eigenvalue is equal or close to zero. Analytically to see, we consider the characteristic polynomial of the matrix, which is given by:

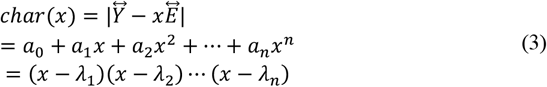

Here, the matrix 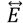denotes the n dimensions identity matrix, *a*_1_ … *a*_*n*_ are the coefficients that represented by the network parameters. Thus, we have

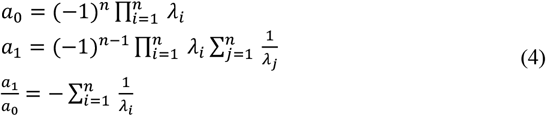

Thus, we consider *a*_1_/*a*_0_ which represents the sum of reciprocals of the eigenvalues. The persist system activity implies that one eigenvalue is equal or close to zero, then the magnitude of the *a*_1_/*a*_0_ would be large. In mathematics, this means two approaches: the first is that *a*_0_ equals to 0 strictly, the second is that *a*_1_ >> *a*_0_.

In our network, the coefficients of the characteristic polynomial *a*_0_ and *a*_1_ can be expressed in terms of the network parameters as:

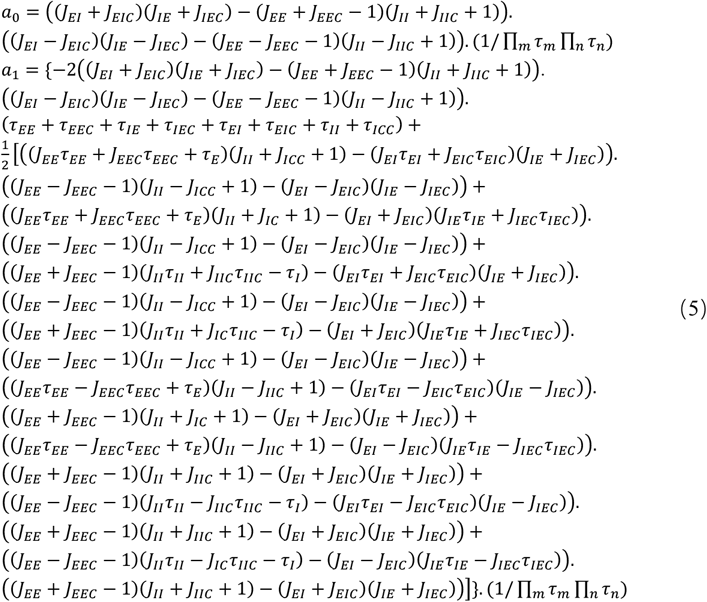

With *m* ∈ {*E, I*}, *n* ∈ {*EE, EEC, IE, IEC, EI, EIC, II, IIC*}

To analyze the expressions above, recognize our assumption that symmetrical network structure leads to symmetrical activities when the system is stable. That means, two sides of the network will be homogeneous with the same firing rate or heterogeneous with “opposite” firing rate. In other words, they will have similar requirements of mathematical expressions for persist activity. In latter competitive condition, due to the firing rate could not be negative, there must be a baseline activity caused by isotropy step-like background input. In the way, the stable activity of four population network is equivalent to the stable activity of two same networks, which contains one E-I couple populations. Furthermore, this application has led to the emergence of two modes. In mode1, inputs from the contralateral side *EEC, EIC, IEC, IIC* act as input in the same side, while in mode 2, inputs from the contralateral side act as inputs in same side, but have the opposite characteristic of excitatory and inhibitory relationship.

Mathematical analyze has proved our assumption above. In fact, the expression *a*_0_ of four population is the product of *a*_0,1_ and *a*_0,2_ which denote the first coefficient of the characteristic polynomial of the two-population network in figure3(a) and figure3(b). In the same way, *a*_1_ is the product of *a*_1,1_ and *a*_1,2_:

**Figure 1.**
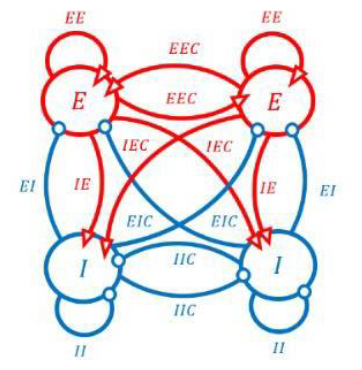
Structure of symmetric competition network with four neural population. All connection strengths and time constants are distributed symmetrically.

**Figure 2.**
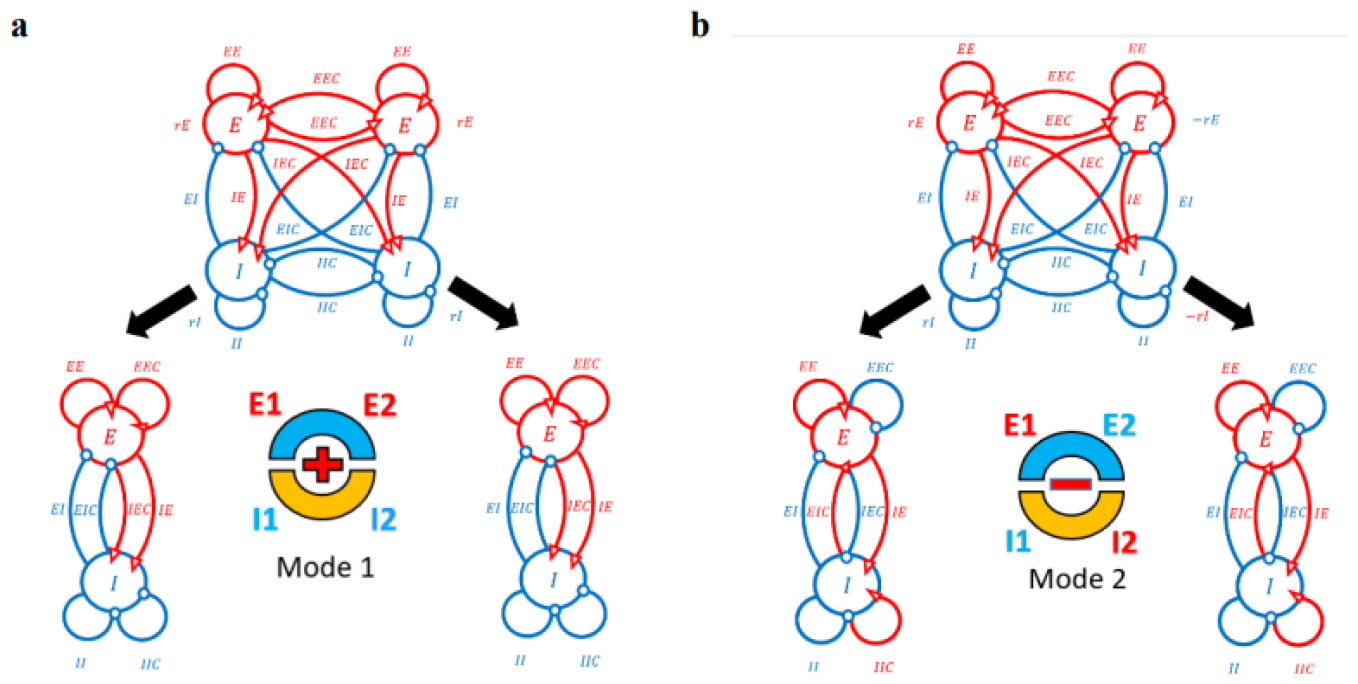
Two Modes of symmetrical activities when the system is stable. (a) Mode1. Two sides of the populations have the same activity while the “cross” connections act as ipsilateral connections. (b) Mode2. Two sides of the populations have the “opposite” activity while the “cross” connections act as ipsilateral connections with reverse relationship of excitatory and inhibitory.

**Figure 3.**
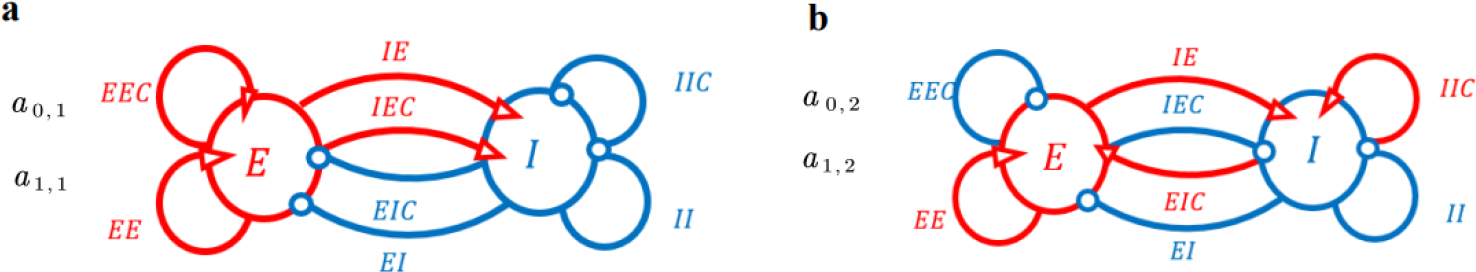
(a) and (b) are networks with of multiple feedback pathways in two conditions.

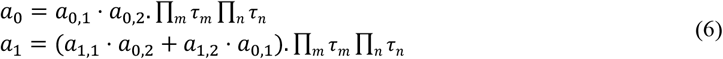

With *m* ∈ {*E, I*}, *n* ∈ {*EE, EEC, IE, IEC, EI, EIC, II, IIC*}.

Expressions of *a*_0,1_, *a*_0,2_, *a*_1,1_, *a*_12_ are

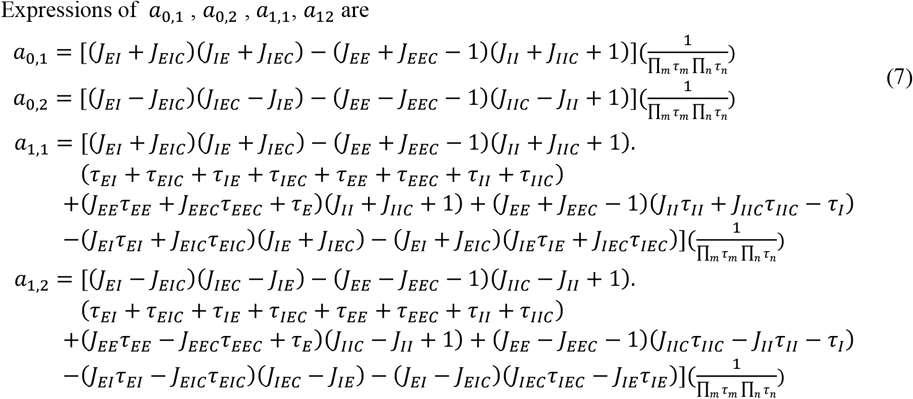

With *m* ∈ {*E, I*}, *n* ∈ {*EE, EEC, IE, IEC, EI, EIC, II, IIC*}

Recall the two scenarios where at least one eigenvalue is close to zero, which means that the characteristic polynomial coefficient *a*_0_ = 0 or *a*_1_ >> *a*_0_. We will discuss the constraints on network parameters and the corresponding network dynamics mechanisms for these two scenarios separately.

## 1. Requirements for *a*_0_ = 0 implies positive feedback

*a*_0_ = 0 quals to *a*_0,1_ = 0 or *a*_0,2_ = 0 or both *a*_0,1_ = 0 and *a*_0,2_ = 0. It is evident that for the vast majority of cases, *a*_0,1_ = 0 and *a*_0,2_ = 0 are generally mutually exclusive, and the conditions under which they are both zero are stringent and thus not count into consideration in our analysis. Instead, we focus on the majority of cases where they are exclusive of each other that *a*_0,1_ = 0 or *a*_0,2_ = 0.

For *a*_0,1_ = 0, we have

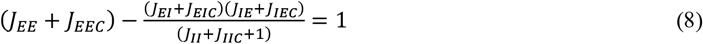

For *a*_0,2_=0, we have

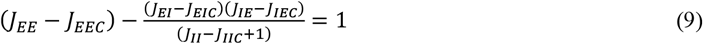

The equilibrium (8) and (9) corresponds that the find-tuning stability condition for mode 1 and mode 2 respectively. The first term in left side represents the overall strength of self-excitation for one population of mode1, and also represents the net self-excitation for one population for mode2 (Fig4A.). The second term represents the strength of self-inhibition throughout the pathways to the inhibitory population. Therefore, for both mode 1 and mode 2, *a*_0_ = 0 means that the difference between the strengths of self-excitation and self-inhibition precisely offsets the leakage losses, thereby allowing the system to maintain sustained activity. This mechanism is known as positive feedback.

**Figure 4A.**
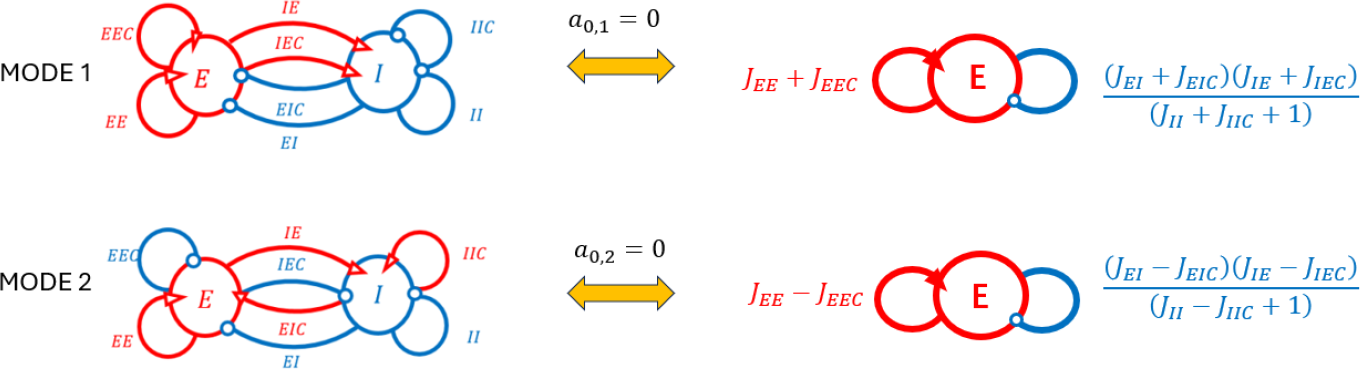
Schematic diagrams of positive feedback under two modes.

**Figure 4B.**
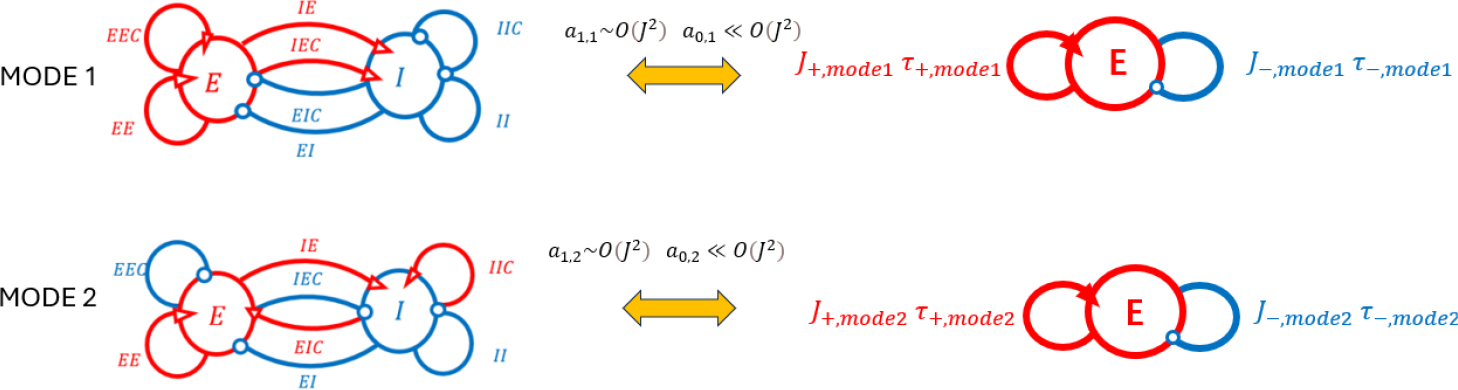
Schematic diagrams of negative derivative feedback under two modes.

## 2. Requirements for *a*_1_ ≫ *a*_0_ implies negative derivative feedback

According to the expressions of *a*_1_ and *a*_0_ in (7), the requirement of *a*_1_ >> *a*_0_ is considered to be two aspects, the time constants *τ* are large or the connection strengths *J* are large but in different order between *a*_1_ and *a*_0_, e.g. *a*_1_∼*O*(*J*^4^) while *a*_0_ ≪ *O*(*J*^4^). However, the large time constant will lead to slow dynamics of the system. Thus we consider the situation about large synaptic strength *J*. Recall from (6), we have

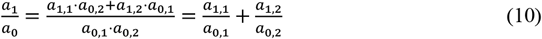

Thus *a*_1_ >> *a*_0_ is mainly derived by *a*_1,1_∼*O*(*J*^2^), *a*_0,1_ ≪ *O*(*J*^2^) or *a*_1,2_∼*O*(*J*^2^), *a*_0,2_ ≪ *O*(*J*^2^) . It’s worth notice that *a*_0,1_ ≪ *O*(*J*^2^) and *a*_0,2_ ≪ *O*(*J*^2^) are mutually exclusive from the expressions (7) when *J*s are large, so *a*_1_/*a*_0_ is dominated by either *a*_1,1_/*a*_0,1_ or *a*_1,2_/*a*_0,2_ and implying that the network is dominated by mode 1 and mode 2 respectively when the network maintains persist activity.

### 2.1 Network dominated by Mode1

Firstly, we consider *a*_1,1_∼*O*(*J*^2^), *a*_0,1_ ≪ *O*(*J*^2^) with large *J*s where the coefficient *a*_1,1_, *a*_0,1_ are dominated by their leading terms with order of *J*^2^ as followings:

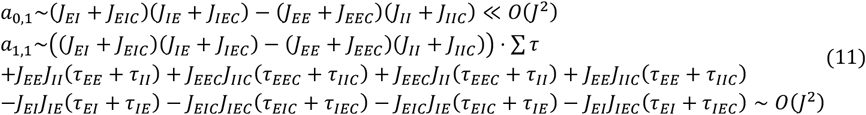

And then the constraints above could lead to

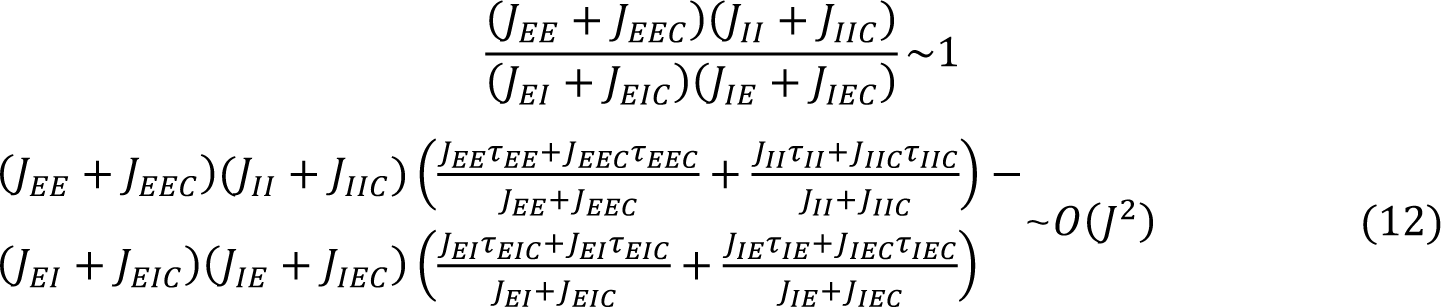

Denote that

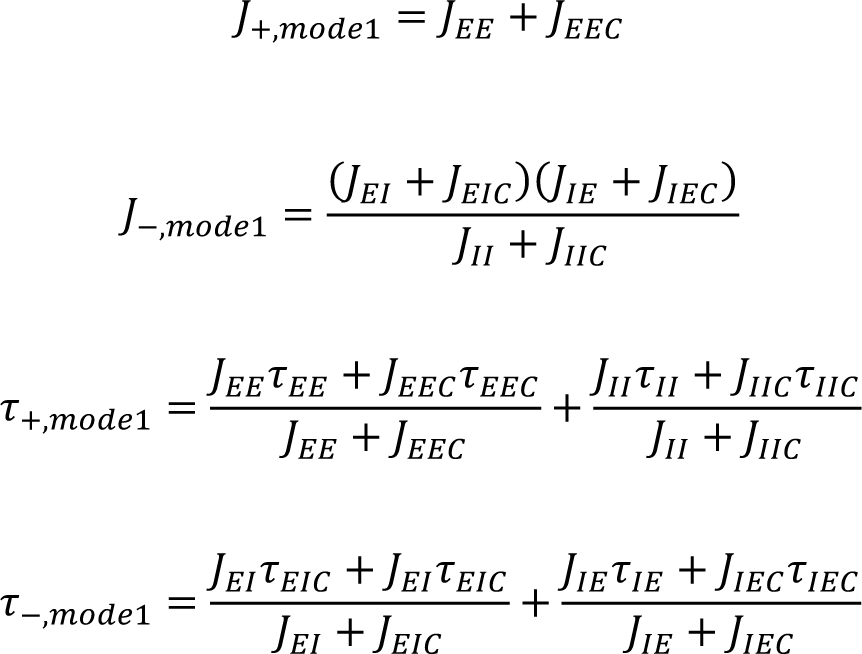

(12) is equivalent to

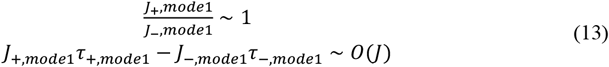

Here, *J*_+,*mode*1_ represents the strength of excitatory feedback and *J*_−_ represent the strength of negative feedback through self-inhibitory population in mode1 (Fig 4B). The term multiplying by the connection strength represents the time constant of positive feedback and negative feedback, which are denoted as *τ*_+,*mode*1_ and *τ*_−,*mode*1_ . The constraints (13) indicate the connection strength of excitatory and inhibitory recurrent activities are balanced, but with different time scale. The property will lead to derivative feedback control according to previous work^[7]^ (Goldman et al., 2013). Simply to see that we consider one neural population with one positive feedback pathway and one negative feedback pathway, which the pathways are equal in strength but different in time constant. Such population is described by

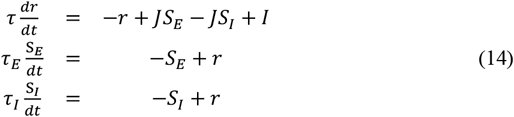

Then, the Laplacian transform of S_*E*_ − S_*I*_ is proportional to

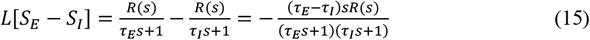

With assumption that the network has persist activity or small changes, the frequency *s* is small then

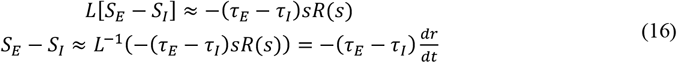

Therefore, derivative feedback arises from the balanced interplay of excitatory and inhibitory strengths across different time scales. The requirement for negative derivative feedback is that *τ*_*E*_> *τ*_*I*_, which is also a requirement for system stability ^[7]^ (Goldman et al., 2013).

In summary, we have identified the conditions under which negative derivative feedback exists when the system is dominated by mode 1. *J*_+,*mode*1_ = *J*_−,*mode*1_ with *τ*_+,*mode*1_ > *τ*_−,*mode*1_.

### 2.2 Network dominated by Mode2

Secondly, we consider *a*_1,2_∼*O*(*J*^2^), *a*_0,2_ ≪ *O*(*J*^2^) with large *J*s where the coefficient *a*_1,2_, *a*_0,2_ are dominated by their leading terms with order of *J*^2^ as followings:

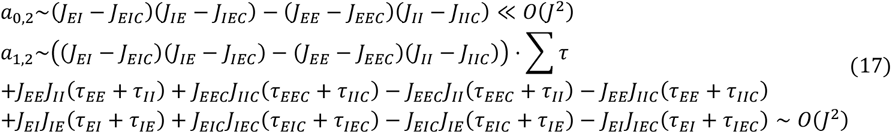

And then the constraints above could lead to

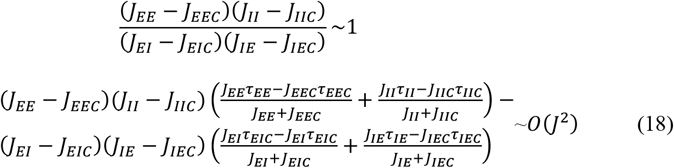

Denote that

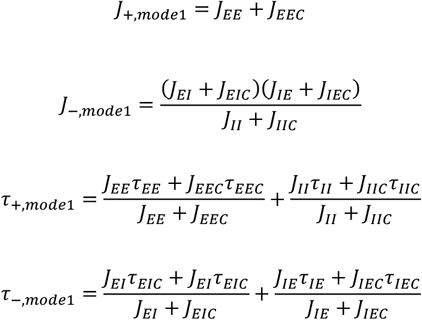

Thus, (18) is equivalent to

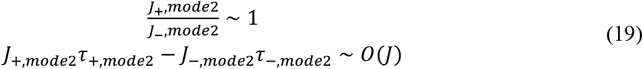

Similarly to mode1, the expressions (19) describes the conditions for derivative feedback when the system is dominated by mode 2, specifically, when negative derivative feedback is present, it should satisfy *τ*_+,*mode*2_ >*τ*_−,*mode*2_.

## 3. Coexist of positive feedback and negative derivative feedback

In our linear system of 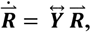, For each right eigenvector 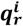and its corresponding eigenvalue **λ**^***i***^ of the matrix 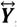, the equation 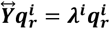 holds true for each ***i*** from 1 to ***n***, where ***n*** denotes the number of state variables. The decay of each mode is exponential, with the time constant 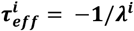. Therefore, the system’s ***τ***_***eff***_ depends on the smallest λ, especially when λ approaches zero. In our analysis, this is equivalent to ***τ***_***eff***_ = ***a***_**1**_/***a***_**0**_ which approximates ***a***_**1**,**1**_/***a***_**0**,**1**_ when mode 1 dominates and ***a***_**1**,**2**_/***a***_**0**,**2**_ when mode 2 dominates.

Recall that the positive feedback strengths in (8) and (9), denote the positive feedback strength in mode 1 and mode 2 as *J*_*pos*,1_ and *J*_*pos*,2_ :

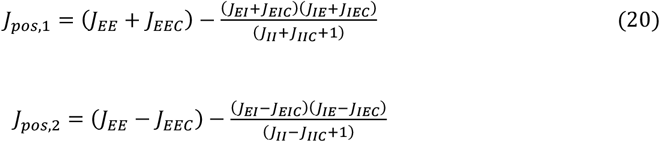

Thus ***a***_**1**,**1**_/***a***_**0**,**1**_ and ***a***_**1**,**2**_/***a***_**0**,**2**_ which represent ***τ***_***eff***_ for the two modes are given by:

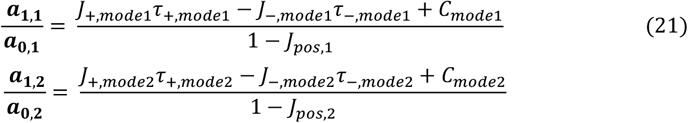

From the previous analysis, it is known that under the condition when *J*s are very large and satisfies *J*_+_ = *J*_−_ and *τ*_+_ > *τ*_−_, the system exhibits negative derivative feedback with intensity *J*_*der*_∼ *J*_+_(*τ*_+_ − *τ*_−_) or *J*_−_(*τ*_+_ − *τ*_−_) (refer to the single population model in (16)). From another aspect, the positive feedback of the system can offset its self-leakage when *J*_*pos*_ = 1. Therefore, the system is able to hold both negative derivative feedback and positive feedback.

## 4. Robustness Against Perturbations

According to previous analysis, negative derivative feedback enables robustness against many perturbations. Here we will analyze the influence of the perturbations in the aspect of changes in the intrinsic neural gains, changes in specific neural sub-population’s gains, changes in excitatory or inhibitory synapses and inactivation of the neurons.

Except that the changes of synapses could be represented by the scaling of the connection strength, other perturbations such as changes in neural gains and inactivation of neurons, could be either represented by the scaling of the connection strength. The reason is that the neural gains of one population is linear to the connections it receives, and the inactivation of one population is negatively linear to all the connections it has. We denote a multiplicative factor *S*_*ij*_, *i, j* ∈ {*E*_1_, *E*_2_, *I*_1_, *I*_2_} to present fluctuations of one synaptic connection strength. After perturbations, the connection strength become *S*_*ij*_*J*_*ij*_.

Firstly, we consider the perturbations that causes symmetric changes of synapses and would not break the symmetric structure of the network. Therefore, the basic form of the coefficients (5) is still available for the new system with perturbations. The simplest condition is that intrinsic neural gains change in the entire network, corresponds to a uniform multiplicative factor *S* to every connection. Under the perturbation, *a*_0_ and *a*_1_ expressions in (5) become 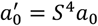 and 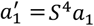 when connection strength *Js* are large and the coefficients *a*_0_ and *a*_1_ are represented by the highest order terms. Because they are scaled similarly, *a*_1_/*a*_0_ will not change for the requirement of negative derivative feedback.

Similarly, changes with excitatory and inhibitory characteristics are symmetrical either, such as inactivation or gains change of E/I populations and synapses. Because the analysis method is similar, we derive a table to represent the three kinds of perturbations in this case. They are changes of E/I neural population gains, changes of E/I connections and inactivation of E/I populations.

The table above shows a robustness against perturbations of symmetrical E/I changes, because the final expression *a*^*′*^ /*a*^*′*^ is same to with large *J*s and the coefficients *a*_0_, *a*_1_ are represented by the highest order terms.

Secondly, we consider the three kinds of perturbations in specific sub-populations: changes in neural gains of one specific sub-population, changes in synapses from one specific sub-population and inactivation of one specific sub-population. Because the change of one specific population has disturbed the symmetrical structure of the network, the *a*_0_ and *a*_1_ expressions in (5) would not make sense at all. Here we derive the expression *a*_0_ of the network without symmetrical structure.

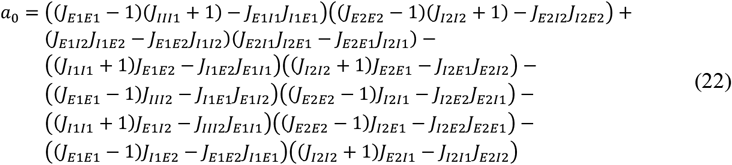

The expression of *a*_1_ is too complex to show in the section, but it is closely related to the expression of *a*_0_, that is, every highest-order item of *a*_0_ and *a*_1_ is a one to one correspondence. The difference is every item in *a*_1_ is multiplicative with a sum of a part of the time constants. Therefore, the conclusion is that if each highest order term of *a*_0_ has a common product of the all connection strength scaling factors Π*S*, then each highest order term of *a*_1_ also has a common product factor Π*S* and in the condition we have *a*_0_^*′*^ = Π*Sa*_0_, *a*_1_^*′*^ = Π*Sa*_1_ when *J* s are large. So, the key point is if each highest term in (20) could be multiplied with a common product of the scaling factors under these three kinds of perturbations. To show influence of different perturbations, we made a table similar to Table1, and take the sub-population *E*1 as an example.

From the expressions above, we conclude that every perturbation of a specific subpopulation could not disturb the form of *a*_1_/*a*_0_ so the network is robust to the perturbations.

## Result

### 1. Persist activity and ramp output

In previous sections, we examined the conditions necessary for negative derivative feedback to occur. We can summarize that such feedback manifests in two distinct firing patterns within a fully symmetric structure: The first mode is characterized by identical firing rates on both sides, while the second mode exhibits inversely related activities against a backdrop of baseline activity. To demonstrate these outcomes, we initially simulate basic firing patterns in response to two types of external input—graded pulse-like and step-like. As depicted in Figure 5 (A) and (C), both firing modes can sustain persistent activity following the pulse-like input, thanks to the negative derivative feedback mechanism. The resulting neural activity maintains a consistent ratio with the graded input. In Figure 5 (B) and (D), the model displays ramp-like activity in reaction to step-like input, with the ramp serving as an integrator of the input over time.

**Figure 5.**
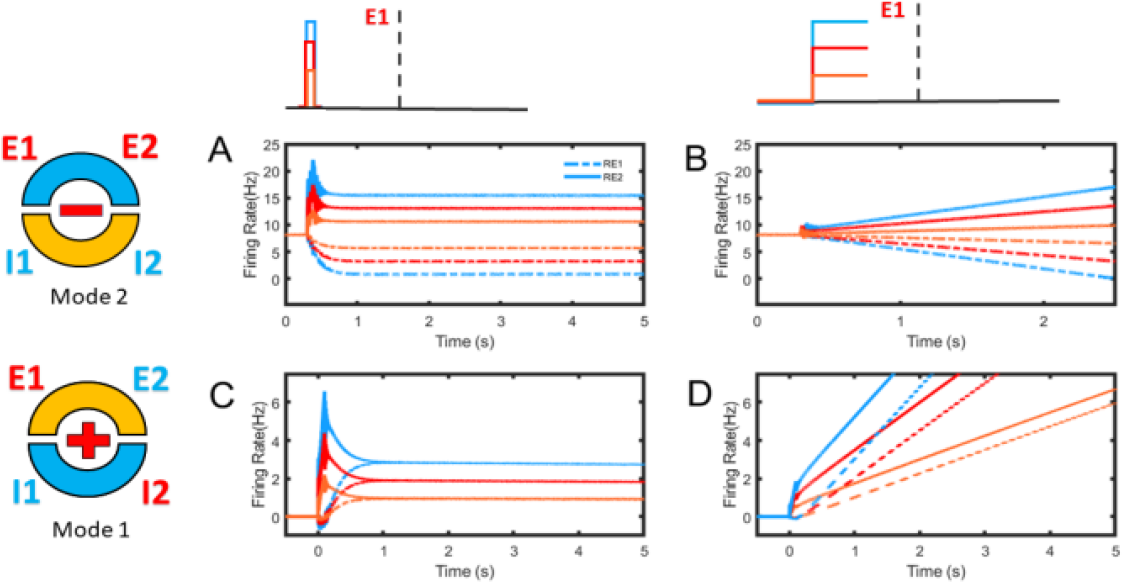
The response patterns of two modes to graded pulse-like and step-like inputs. The modes are represented as a ring on the left, with the plus and minus symbols indicating the reciprocal excitation and inhibition relationships between the two halves. The shapes of the external inputs are depicted at the top. In (A) and (C), both sides are able to sustain ongoing neural activity post-input. Upon stabilization, mode 1 exhibits identical activity levels on both sides, while mode 2 displays “inverse” activities against the baseline background activity. Before achieving this stability, however, the two modes exhibit asymmetry: they are either opposite or identical. Panels (B) and (D) show ramp-like activity induced by step-like input, demonstrating the temporal integration of the input by the ramp. This integration is subject to an asymmetric bias due to the continuous influence of the asymmetric external input.

### 2. Symmetric activity and asymmetric bias

Symmetry is a crucial characteristic of the networks we study. It manifests as either complete alignment or complete opposition relative to the baseline background activity, aligning with the two firing modes discussed in previous sections. Typically, a symmetrical network structure promotes symmetrical activity independent of external input. As such, the network’s dynamic characteristics are determined solely by internal structural parameters like connection weights and time constants.

Specifically, when exposed to asymmetric external inputs, the network exhibits asymmetric activity during the input period. This asymmetry persists until the input decays to zero or is removed, after which the system regains stability and symmetry, as illustrated in Figures 5 (A, C) and 6 (A, D). Conversely, if the external input is symmetrical, the network’s activity remains symmetrical, regardless of whether it reaches stability (see Figures 6 B, C, E, F). This consistency occurs because both key determinants of the system’s dynamics—the external input and the internal structure—are symmetrical. Moreover, symmetric external input can result in two distinct patterns: either complete similarity or complete opposition.

**Figure 6.**
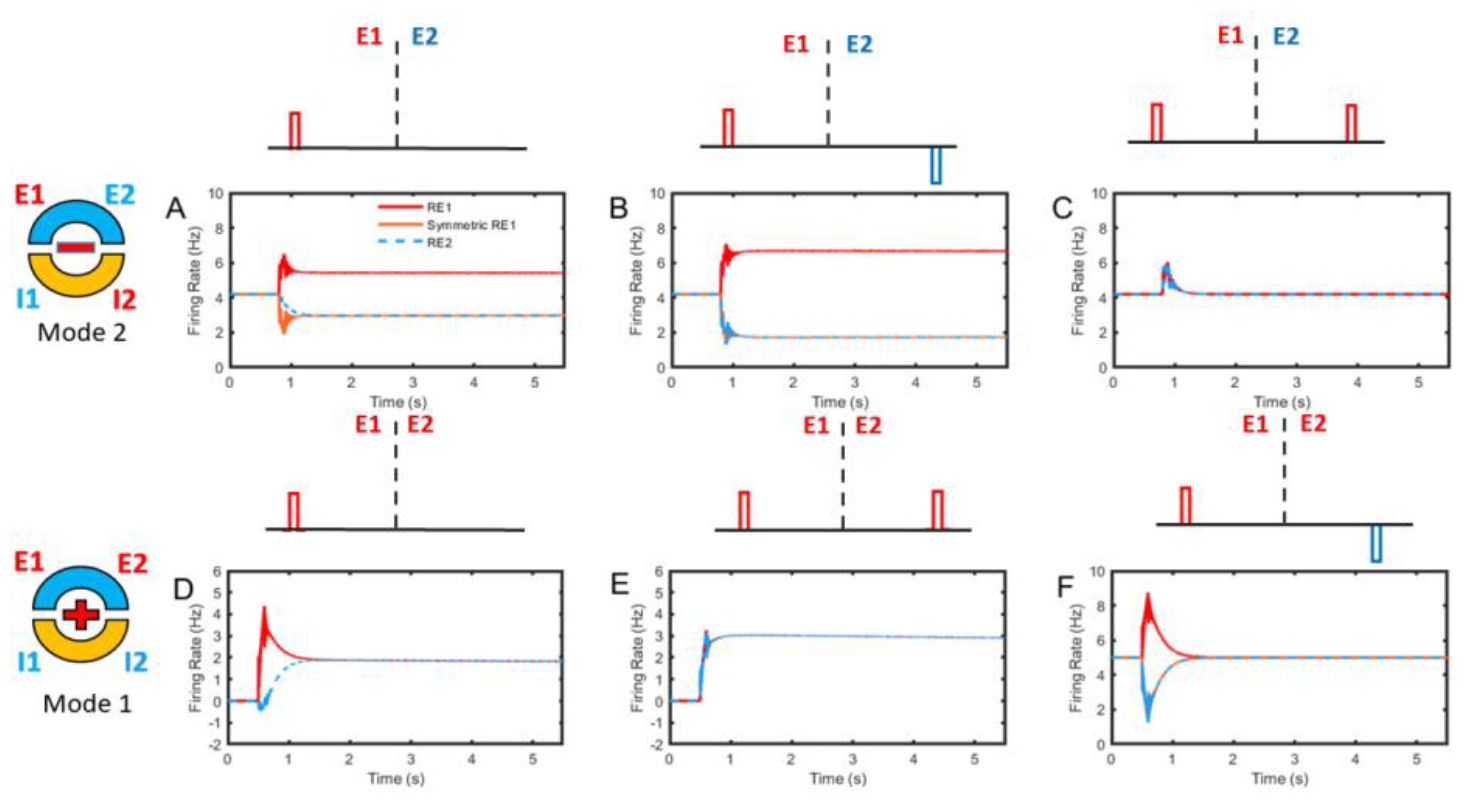
The reaction activity of two modes responds to the symmetrical pulse-like external inputs of two patterns. A, D: The activity caused by unilateral external input. The network has asymmetric activity during the action time of the asymmetric external input. B, E: External input is consistent with the intrinsic structure of two modes. Effect of each unilateral input is same so the overall activity is sum to be double as the activity caused by unilateral input of each side. C, F: Each external unilateral input causes opposite activity, and finally offset each other.

Here, we explore how two activity modes react to symmetric pulse-like inputs. In mode 2, where each side inhibits the other equally, the impact of a positive unilateral input to E1 is identical to a negative unilateral input to E2. Consequently, the overall effect of symmetrical opposite external inputs is twice that of each unilateral input (see Fig 6. A, B). However, if the external inputs are identically symmetric, the effects of the two unilateral inputs cancel each other out due to the equivalent mutual inhibition, resulting in zero net activity (see Fig 6.C).

Similarly, in mode 1, where both sides excite each other, a symmetrically identical input will amplify the effect of a unilateral input (see Fig 6. D, E). Conversely, symmetrical opposite inputs will neutralize each other due to equivalent mutual excitation (see Fig 6.F).

We next examine how two modes respond to two symmetric step-like inputs. In mode 2, due to equivalent mutual inhibition, the ramp output from a positive unilateral step input to E1 matches the effect of a negative unilateral step input to E2. Symmetric opposite step-like inputs, aligned with the network’s intrinsic structure, effectively double the slope of the ramp activity (see Fig 7.A, B). Conversely, if the step inputs are symmetrically identical, they produce no increase in activity, as identical unilateral step inputs generate opposite slopes that cancel each other out (see Fig 7.C). It’s important to note that the cancellation of increasing activity does not imply a decay to zero; the activity preceding the cancellation is maintained. The question then arises: what is the source of this residual activity? As previously demonstrated, the ramps of RE1 and RE2 caused by unilateral step inputs are asymmetrical; they have opposite slopes but originate from different states, leading to an asymmetrical bias (see Fig 6.B, Fig 7.A). The offset occurs only in the slopes, not in the activities arising from different original states.

**Figure 7.**
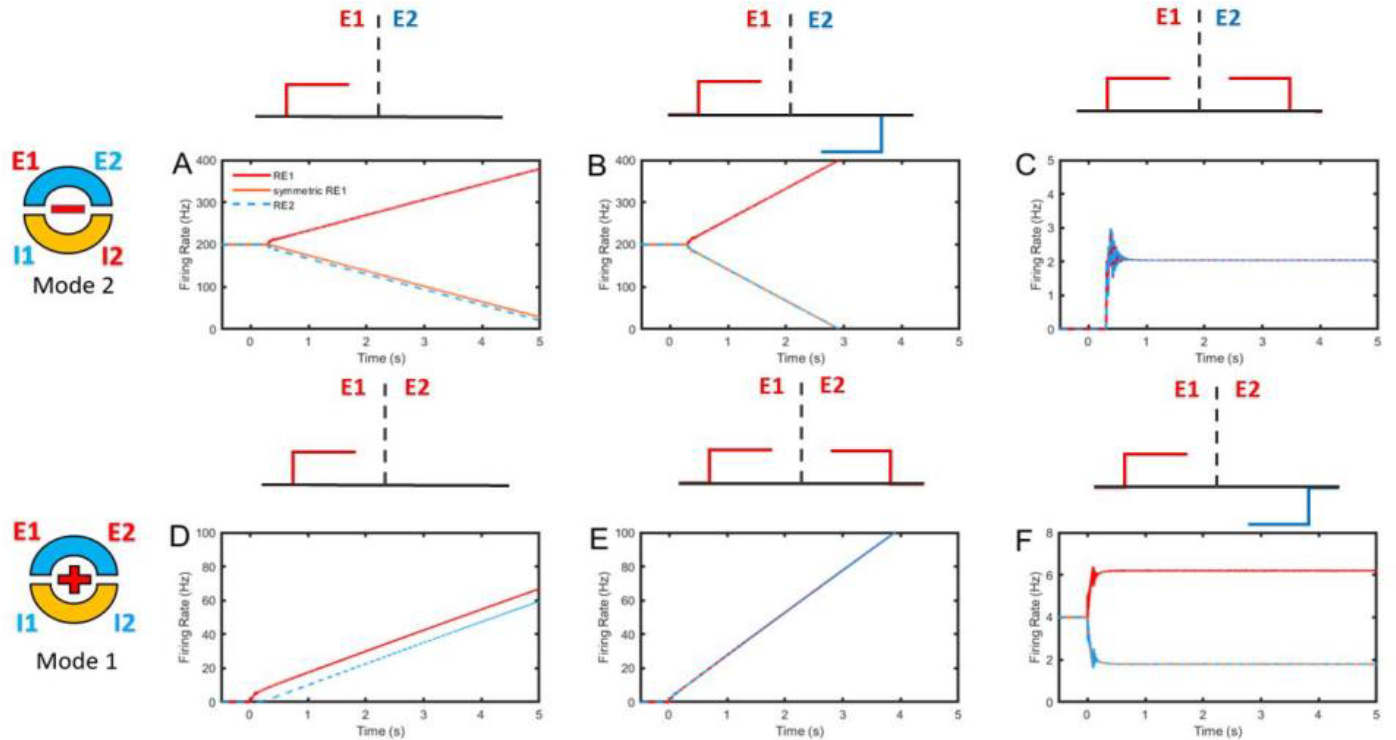
How two modes react to symmetrical step-like external inputs of two patterns. Panels A and D show the activity triggered by unilateral external inputs, where the network displays asymmetric activity throughout due to the continuous action of asymmetrical step-like inputs. Panels B and E depict inputs that align with the intrinsic structure of the two modes, where the overall activity’s slope is twice that of the activity induced by each unilateral step input. In Panels C and F, each unilateral external input creates an opposing slope on the other side, ultimately cancelling each other out. As a result, there is no overall increase in activity, but the initial activity that occurs before the mutual reaction and cancellation is preserved and maintained.

Physically speaking, each synapse is characterized by its time constant and delay. When identical symmetrical inputs are administered to E1 and E2, they instantaneously trigger the same activity. Initially, mutual inhibition does not occur due to synaptic time delays. Subsequently, as mutual inhibition takes effect, the increasing activities counterbalance each other due to their opposing slopes, yet the activities preceding mutual inhibition are preserved and maintained. This persistent activity in mode 2 mirrors the persistent activity in mode 1, which is elicited by symmetrical, same pulse-like inputs (see Fig 6.E). This mechanism also underlies the background activity in competitive networks.

In mode 1, where there is mutual excitation between both sides, a positive symmetrical step input will double the slope (see Fig 7. D, E). When symmetrical opposite step-like inputs are applied, each unilateral step input generates ramp activities with opposing slopes. These activities are initially offset by mutual excitation, yet any initial opposing activity prior to the onset of mutual excitation is preserved (see Fig 7.F).

### 3. Difference in firing pattern compared with two-population model

Our analysis of the network incorporates a significant dimension reduction. When the system is stable, the network activity in both modes resembles that of a dual-network system, each consisting of two populations. Mathematical analysis has demonstrated the correlation between the four-population model and the two-population model, indicating that the prerequisites (*a*_1_/*a*_0_) for persistent activity are the same in both modes. To highlight this equivalence and correlation, we compare the firing patterns of the two models. The parameters of the two-population model are derived from the dimension reduction applied to the four-population model in each mode separately.

The results reveal that the persistent activity in the two networks across both modes is equivalent (Fig 8. A, B). “Equivalence” here implies that both models exhibit the same firing rates and synaptic inputs, but only in a stable state. In other words, for persistent activities, if the firing rates RE1 for the four-population model and rE for the two-population model are set to the same value, then they will share identical firing rates for inhibitory populations RI1 and rI, as well as the same synaptic inputs SEE and sEE. However, this equivalence in persistent activity does not mean that the states are identical at all times. They originate from different unstable states when triggered by the same external inputs and will lead to different firing patterns once stabilized. Conversely, if the two models exhibit the same firing patterns when stable, it does not necessarily mean that the external inputs to both models are identical.

**Figure 8.**
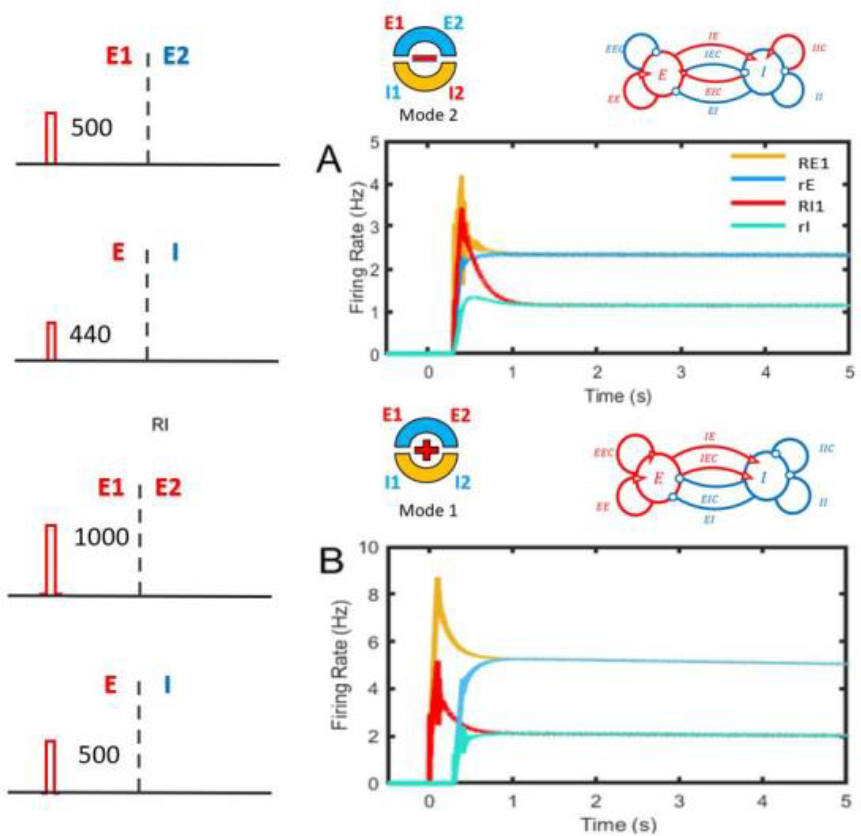
The comparison of four population model and two population model. They have equivalence firing rates and synaptic inputs but different original unstable states and triggered by different external inputs. A. Two model’s firing rate of mode2. B. Two model’s firing rate of mode1.

### 4. Roubtness and perturbations

We have analyzed why negative derivative feedback could be robust against such perturbations. In this section, we will test the model with symmetric perturbations as outlined in Table 1, demonstrating the differences in robustness between positive feedback, which lacks resilience, and negative derivative feedback. To make the simulation results more intuitive, we use mode 2 to illustrate the effects of perturbations and robustness. The figures display the activity of E1 and E2 both without changes (in blue) and with perturbations (in orange). Initially, we address changes in intrinsic neural gains where all populations receive either slightly more or less synaptic input. Setting the perturbation at a 5% increase, the results reveal that positive feedback fails to maintain persistent activity and is excessively strengthened (Fig 9 A’). If the initial balance between positive and negative feedback strengths is 1, this balance is disrupted as both are increased by the same multiple. However, with negative derivative feedback, where positive and negative feedback are balanced in strength, this relationship remains unchanged.

**Table 1.** Analysis of the symmetrical perturbations with E/I characteristic.

| Perturbation Type | Perturbation Expression | Explanation of the expression | Expression of $a_0'$ and $a_1'$ | Expression of $a_1'/a_0'$ |
| --- | --- | --- | --- | --- |
| Changes of E/I neural gains | $S_E J_{Em}, S_E J_{EmC}$<br>$S_I J_{In}, S_I J_{InC}$,<br>$m, n = E, I$ | $S_E J_{Em}$ and $S_I J_{In}$ denote connections that E or I populations gained from other populations under the perturbation. | $a_0' = S_I^2 S_E^2 a_0$<br>$a_1' = S_I^2 S_E^2 a_1$<br>With large $J$s | $\frac{a_1'}{a_0'} = \frac{a_1}{a_0}$ |
| Changes of E/I synapses | $S_E J_{mE}, S_E J_{mEC}$<br>$S_I J_{nI}, S_I J_{nIC}$,<br>$m, n = E, I$ | $S_E J_{mE}$ and $S_I J_{nI}$ denote the E or I synapses that originated from E or I populations under the perturbation. | $a_0' = S_I^2 S_E^2 a_0$<br>$a_1' = S_I^2 S_E^2 a_1$<br>With large $J$s | $\frac{a_1'}{a_0'} = \frac{a_1}{a_0}$ |
| Inactivation of E/I populations | $S_E^2 J_{EE}, S_E^2 J_{EEC}$<br>$S_I^2 J_{II}, S_I^2 J_{IIC}$<br>$S_E S_I J_p$<br>$p = EI, IE, EIC, IEC$ | In inactivation perturbations, neural gains and E/I synapses has changed together, so the square or multiple scaling factor shows the two aspects of the changes: neural gains and synapses. | $a_0' = S_I^4 S_E^4 a_0$<br>$a_1' = S_I^4 S_E^4 a_1$<br>With large $J$s | $\frac{a_1'}{a_0'} = \frac{a_1}{a_0}$ |

**Table 2.**
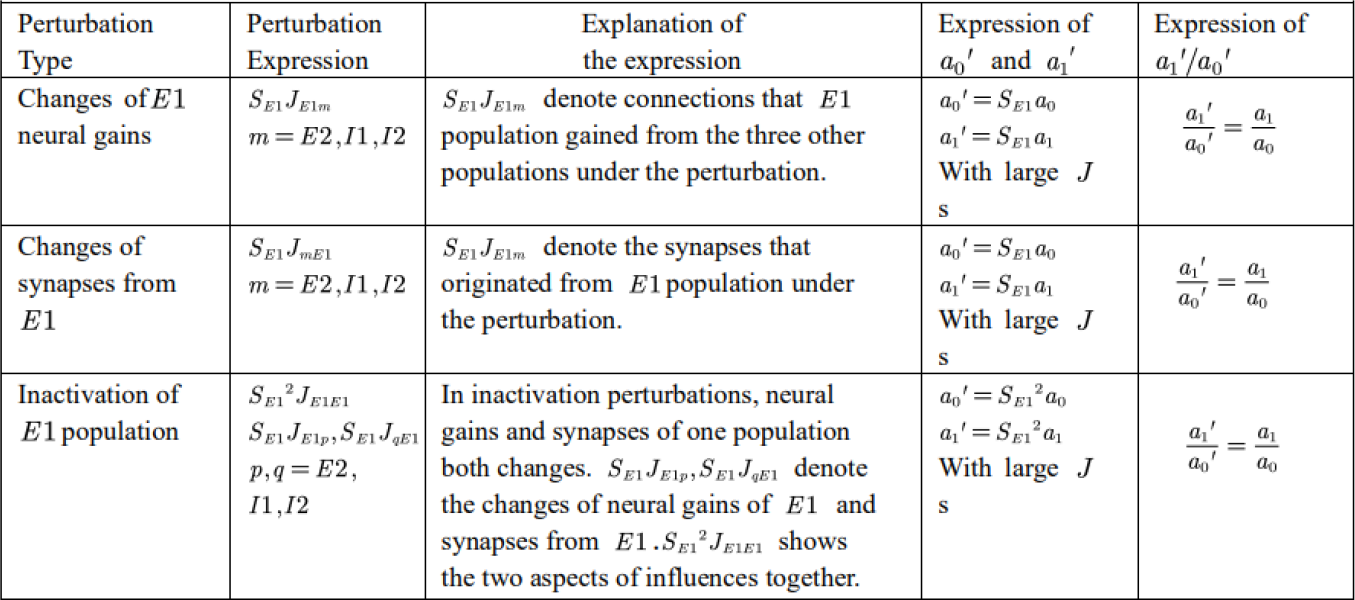
Analysis of the perturbations of a specific population. (Take the population *E*l as an example).

**Figure 9.**
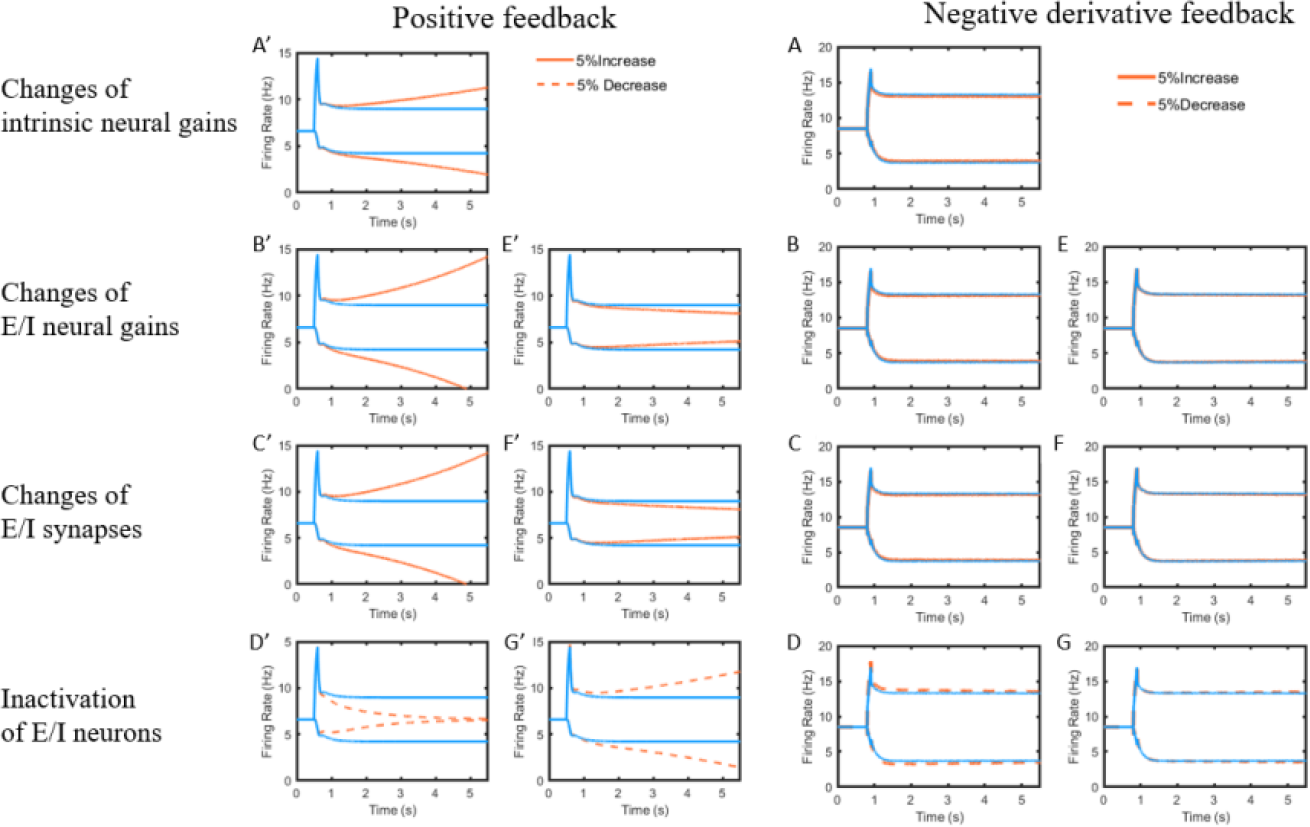
The reaction of positive feedback and negative feedback model to the symmetric perturbations.

Secondly, we evaluate changes in neural gains and synaptic adjustments for excitatory and inhibitory populations, with excitatory changes detailed in the left column and inhibitory changes in the right. These two types of perturbations share similar mathematical expressions and, consequently, impart identical effects on the activities (Figure 9. B’, C’, E’, F’). Evidently, the positive feedback model lacks robustness; neither excitatory nor inhibitory changes can preserve the fine tuning. In contrast, the negative derivative feedback model maintains its robustness against these perturbations, as the balance condition (referenced in the Method section) remains unchanged.

Finally, we examine the inactivation of excitatory or inhibitory populations. In the positive feedback model, such inactivation significantly disrupts fine tuning and induces perturbations. Conversely, in the negative feedback model, the activity remains stable, demonstrating its robustness.

The asymmetric perturbation is mainly about changes of sub-population. According to mathematical analysis in Method, perturbations of one subpopulation has not changed basic *a*_1_/*a*_0_ expression for negative derivative feedback, but has changed *a*_0_and disturbed *a*_0_ = 0 for positive feedback.

To show the robustness and perturbations, we simulate the activity under the changes of E1 and I1 neural gains, changes of synapses of E1 and I1 and inactivation of E1 and I1. All results shows that negative derivative feedback has robustness against these perturbations, but positive feedback is disturbed by these perturbations and could not maintain persist activity again (Fig 10).

**Figure 10.**
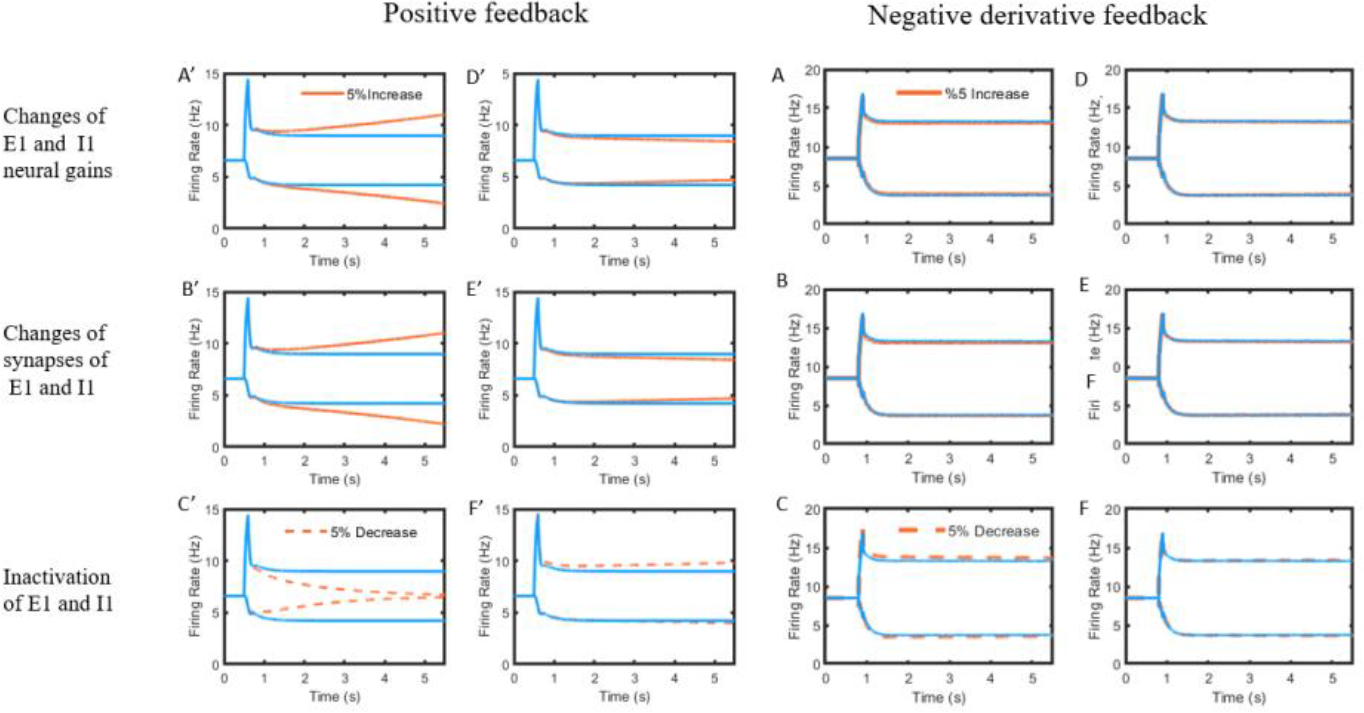
The reaction of positive feedback and negative feedback model to the asymmetric perturbations of one specific sub-population.

Our results reveal a novel competitive neural network architecture consisting of two symmetrical EI populations, with all populations fully interconnected. This structure not only supports the robustness of negative derivative feedback against common perturbations but also offers greater biological plausibility.

A key assumption of our network is that in the absence of external inputs, the symmetrical structure will result in symmetrical activities. This is because the network’s final state is determined by its structural parameters. This assumption facilitates a dimensional reduction perspective, suggesting that a four-population network exhibiting persistent activity is effectively equivalent to two identical two-population networks on both sides.

Mathematical analysis further indicates that this dimensional reduction leads to two distinct activity modes: mode 1, where neural activities on both sides are symmetrically identical, and mode 2, where activities are symmetrically opposite. Since firing rates cannot be negative, “opposite” here implies that the activity includes a baseline that is not zero. Figure 7.C illustrates that for mode 2, a shared positive step input on both sides can establish an activity baseline. The transition between these two modes depends on the balance of excitation and inhibition between the two sides. In mode 1, mutual activation on both sides exceeds mutual inhibition, whereas in mode 2, mutual inhibition dominates over mutual activation.

As a verification, the simulation shows the equivalence in persist activity of four population model and two population model (Fig.8). They have equivalence firing rates and synaptic inputs but different original unstable states. Therefore, to trigger the equivalent persist activity of two models, they need different external inputs.

The symmetry dimension reduction can provide us a simplification to analyze negative derivative feedback because two-population networks and four-population networks have the same requirements for negative derivative feedback. To illustrate the requirements in specific, negative derivative feedback needs balance in positive and negative feedback strengths, but lager arrival time for positive feedback than negative feedback with the form of *J*_+_ = *J*_−,_, *τ*_+_ > *τ*_−_ which correspond the traditional model that one direction (from and to which population) just has one connection. The simplification has also suggested us a perspective of synapse integration that multiple connections in one direction could be integrated as one. The integrated connection strength is the sum of the original connections, and the time constant of the integrated connection is the average of original time constants with a strength weighted scaling factor. Therefore, we could extend the four-population model with multiple connections between two specific sub-populations.

One of the characteristics of negative derivative feedback is it has robustness to many common perturbations. In nerve system, common perturbations are changes of neural gains, changes of synapses and inactivation of neurons. Combined with the model structure, we divide these perturbations into symmetric perturbations caused by changes of E/I population on both sides and asymmetric perturbations caused by changes of one sub-population. All the perturbations could be mathematically represented by a multiplicative scaling factor to connection strengths. Analyze shows these perturbations could not changes the basic requirements for negative derivative feedback, simulations also show the strong robustness to these perturbations as verification. However, by comparison, positive feedback has no robustness to any kind of perturbations.

External inputs have many influences to activities. The persist activity is proportional to graded pulse-like external input, and the slope of ramp-like activity is proportional to graded step-like input. If the external input is asymmetric, the activity of the system is also asymmetric during its action. The asymmetric bias is clear when the network is activated by step-like unilateral external input (Fig.7 A, D). If the external input is symmetric, the activity will always be symmetric. What’s more, the symmetric same external input will double the activity caused by unilateral input in mode 1 because mutual activation, but in mode2, because of the equivalent mutual inhibition, the effect of each unilateral input will offset each other. In contrast, the symmetric opposite external input will double the activity caused by unilateral input in mode 2 it is consistent with the mutual inhibition structure, but effect of each unilateral input will offset, because of the equivalent mutual activation will weaken the difference between two side.

Symmetry is important for competition network. In addition to the symmetric model, the question is, what’s the requirements for negative derivative feedback of asymmetric four-population model? Because it could not be dimensionally reduced, expression (22) shows the restrictions to find positive feedback pathway and negative feedback pathways on a global perspective. When there are more than two populations, positive feedback and negative feedback pathways are not well defined, so we could not draw an intuitive conclusion. Positive feedback is used in many traditional competition networks to maintain the activity, such as working memory and neural integrator. Our model is still applicable to traditional positive feedback models by symmetric dimensional reduction, and still has more biological rationality at this level. Besides, for some cases, negative derivative feedback may not be applicable. The model has a wide range of practicability, but how to apply the model to experiments and explain physiological recordings are still needs further exploration.

